# Alpha-frequency feedback to early visual cortex orchestrates coherent natural vision

**DOI:** 10.1101/2023.02.10.527986

**Authors:** Lixiang Chen, Radoslaw Martin Cichy, Daniel Kaiser

## Abstract

During natural vision, the brain generates coherent percepts by integrating sensory inputs scattered across the visual field. Here, we asked whether this integration process is mediated by rhythmic cortical feedback. In EEG and fMRI experiments, we experimentally manipulated the demand for integration by changing the spatiotemporal coherence of natural videos presented across visual hemifields. Our EEG data revealed that information about incongruent videos is coded in feedforward-related gamma activity while information about congruent videos is coded in feedback-related alpha activity, indicating that integration is indeed mediated by rhythmic feedback. Our fMRI data identified scene-selective cortex as a likely source of this feedback. Analytically combining our EEG and fMRI data further revealed that feedback-related representations in the alpha band shape the earliest stages of visual processing in cortex. Together, our findings indicate that the construction of coherent visual experiences relies on rhythmic cortical feedback that fully traverses the visual hierarchy.

## Introduction

We consciously experience our visual surroundings as a coherent whole that is phenomenally unified across space ^1,2^. In our visual system, however, inputs are initially transformed into a spatially fragmented mosaic of local signals that lacks integration. How does the brain integrate this fragmented information across the subsequent visual processing cascade to mediate unified perception?

Classic hierarchical theories of vision posit that integration is solved during feedforward processing ^3,4^. On this view, integration is hard-wired into the visual system: Local representations of specific features are integrated into more global representations of meaningful visual contents through hierarchical convergence over features distributed across visual space.

More recent theories instead posit that visual integration is achieved through complex interactions between feedforward information flow and dynamic top-down feedback ^5–7^. On this view, feedback information flow from downstream adaptively guides the integration of visual information in upstream regions. Such a conceptualization is anatomically plausible, as well as behaviorally adaptive, as higher-order regions can flexibly adjust current integration demands through the visual system’s abundant top-down connections ^8–10^.

However, the proposed interactions between feedforward and feedback information pose a critical challenge: Feedforward and feedback information needs to be multiplexed across the visual hierarchy to avoid unwanted interferences through spurious interactions of these signals. Previous studies propose that neural systems meet this challenge by routing feedforward and feedback information in different neural frequency channels: high-frequency gamma rhythms may mediate feedforward propagation, whereas low-frequency alpha and beta rhythms carry predictive feedback to upstream areas ^11–14^.

Here, we set out to test the hypothesis that rhythmic coding acts as a mechanism mediating coherent visual perception. We used a novel experimental paradigm that manipulated the demand for spatial integration through the spatiotemporal coherence of natural videos shown in the two visual hemifields. Combining EEG and fMRI measurements, we show that when inputs are integrated into a coherent percept, cortical alpha dynamics carry stimulus-specific feedback from high-level visual cortex to early visual cortex. Our results show that spatial integration of natural visual inputs is mediated by feedback dynamics that traverse the visual hierarchy in low-frequency alpha rhythms.

## Results

We experimentally mimicked the spatially distributed nature of natural inputs by presenting eight 3-second natural videos (Fig. 1A) through two circular apertures right and left of fixation (diameter: 6° visual angle, minimal distance to fixation: 2.64°). To assess spatial integration in a controlled way, we varied how the videos were presented through these apertures (Fig. 1B): In the right- or left-only condition, the video was shown only through one of the apertures, providing a baseline for processing inputs from one hemifield, without the need for spatial integration across hemifields. In the congruent condition, the same original video was shown through both apertures. Here, the input had the spatiotemporal statistics of a unified scene expected in the real world and could thus be readily integrated into a coherent unitary percept. In the incongruent condition, by contrast, the videos shown through the two apertures stemmed from two different videos. Here, the input did not have the spatiotemporal real-world statistics of a unified scene, and thus could not be readily integrated. Contrasting brain activity for the congruent and incongruent condition thus reveals neural signatures of spatial integration into unified percepts across visual space.

**Fig. 1.**
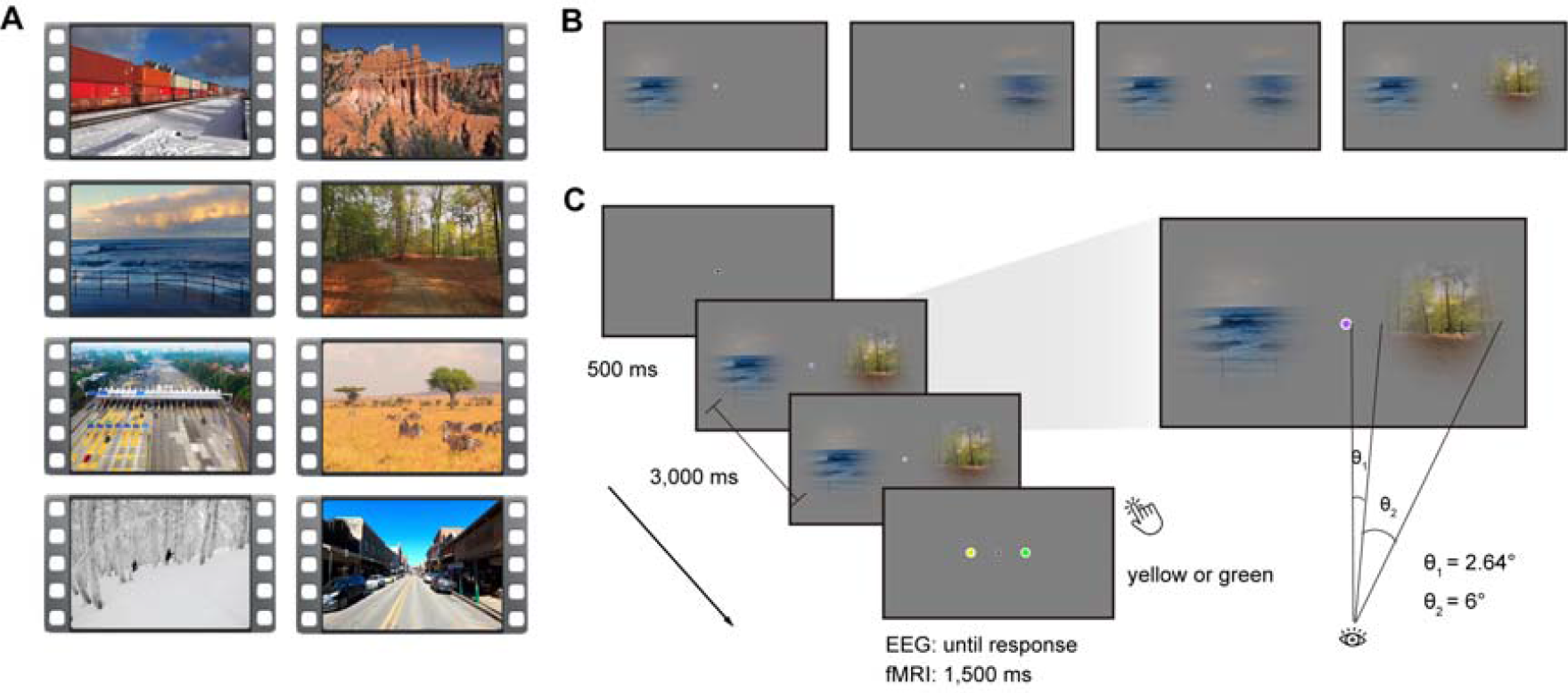
Stimuli and Experimental design. **A)** Snapshots from the eight videos used. **B)** In the experiment, videos were either presented through one aperture in the right or left visual field, or through both apertures in a consistent or inconsistent way. **C)** During the video presentation, the color of the fixation dot changed periodically (every 200 ms). Participants reported whether a green or yellow fixation dot was included in the sequence.

Participants viewed the video stimuli in separate EEG (n = 48) and fMRI (n = 36) recording sessions. Participants performed an unrelated central task (Fig. 1C) to ensure fixation and to allow us to probe integration processes in the absence of explicit task demands.

Harnessing the complementary frequency resolution and spatial resolution of our EEG and fMRI recordings, we then delineated how inputs that either do or do not demand integration into a coherent percept are represented in rhythmic neural activity and regional activity across the visual hierarchy. Specifically, we decoded between the eight different videos in each of the four conditions from frequency-resolved EEG sensor patterns ^15,16^ and from spatially resolved fMRI multi-voxel patterns ^17^.

Our first key analysis determined how the feedforward and feedback information flows involved in processing and integrating visual information across space are multiplexed in rhythmic codes. We hypothesized that conditions not affording integration lead to neural coding in feedforward-related gamma activity ^11,14^, whereas conditions demanding spatiotemporal integration lead to coding in feedback-related alpha/beta activity ^11,14^.

To test this hypothesis, we decoded the video stimuli from spectrally resolved EEG signals (aggregated within the alpha, beta, and gamma frequency bands) during the whole stimulus duration (Fig. 2A; see Methods for details). Our findings supported our hypothesis. We observed that incongruent video stimuli, as well as single video stimuli, were decodable only from the gamma frequency band (all *t*(47) > 3.41, *p* < 0.001; Fig. 2B and 2C). By stark contrast, congruent video stimuli were decodable only from the alpha frequency band (*t*(47) = 5.43, *p* < 0.001; Fig. 2C). Comparing the pattern of decoding performance across frequency bands revealed that incongruent video stimuli were better decodable than congruent stimuli from gamma responses (*t*(47) = 3.04, *p* = 0.004) and congruent stimuli were better decoded than incongruent stimuli from alpha responses (*t*(47) = 2.32, *p* = 0.025; interaction: *F*(2, 94) = 7.47, *p* < 0.001; Fig. 2C). The observed effects also held when analyzing the data continuously across frequency space rather than aggregated in pre-defined frequency bands (see Fig. S2), and were not found trivially in the evoked broadband responses (see Fig. S3). Together, our results demonstrate multiplexing of visual information in rhythmic information flows. When no integration across hemifields was required, visual feedforward activity is carried by gamma rhythms. When spatiotemporally coherent inputs demanded integration, integration-related feedback activity is carried by alpha rhythms.

**Fig. 2.**
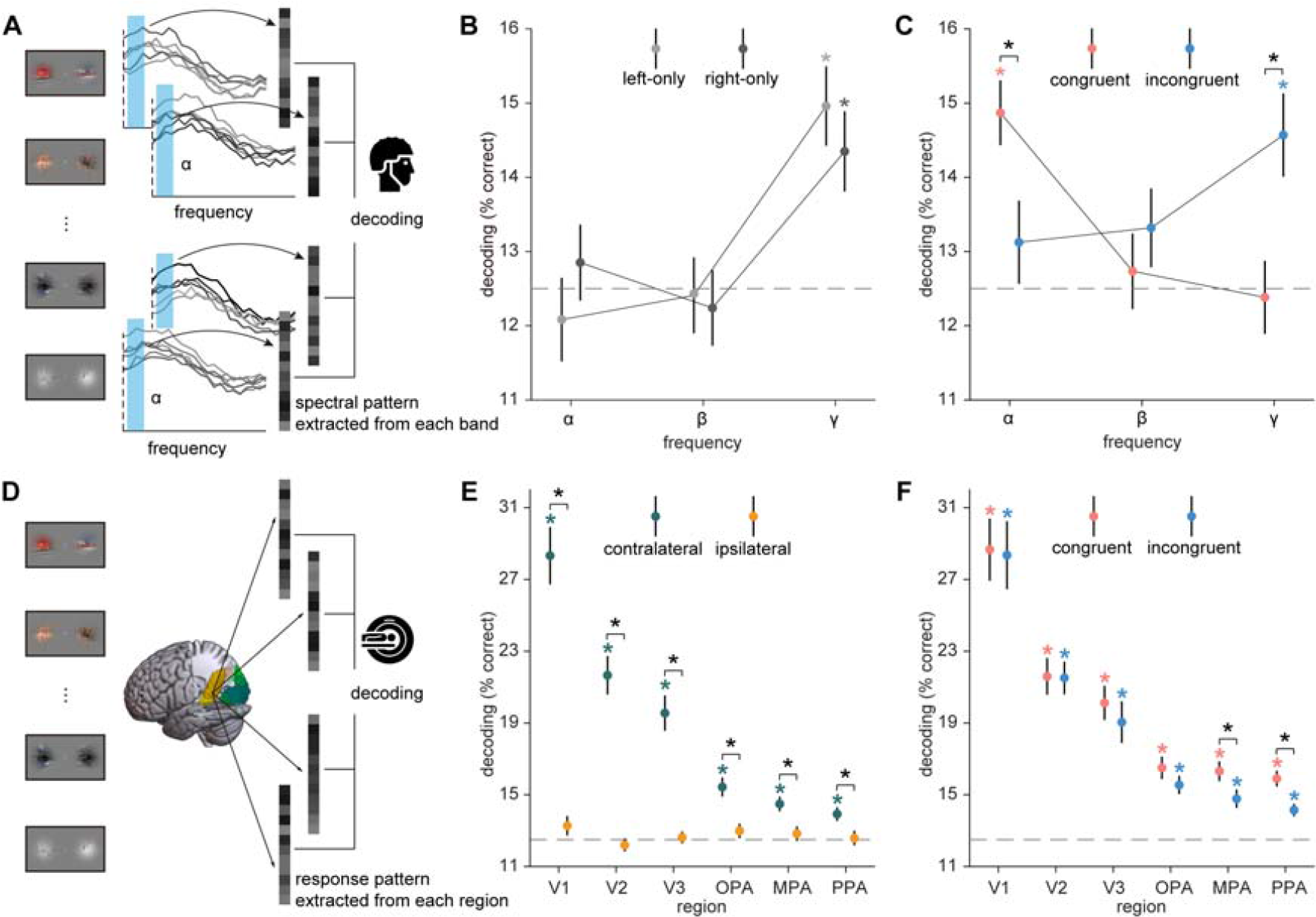
EEG and fMRI decoding analysis. **A)** Frequency-resolved EEG decoding analysis. In each condition, we used eight-way decoding to classify the video stimuli from patterns of spectral EEG power across electrodes, separately for each frequency band (α, β, γ). **B, C)** Results of EEG frequency-resolved decoding analysis. The incongruent and single video stimuli were decodable from gamma responses whereas congruent stimuli were decodable from alpha responses, suggesting a switch from dominant feedforward processing to the recruitment of cortical feedback. **D)** fMRI decoding analysis. In each condition, we used eight-way decoding to classify the video stimuli on response patterns in each region of interest (V1, V2, V3, OPA, MPA, PPA). **E)** Results of fMRI decoding analysis for the right- and left-only conditions. Single video stimuli were decodable in regions contralateral to the stimulation, but not in ipsilateral regions. **F)** Results of fMRI decoding analysis for the congruent and incongruent conditions. Video stimuli were decodable in both conditions in each of the six regions. Congruent stimuli were decoded better than incongruent stimuli in scene-selective cortex (MPA and PPA), suggesting that these regions integrate information across visual hemifields. Error bars represent standard errors. *: *p* < 0.05 (FDR-corrected).

The observation of a frequency-specific channel for feedback information underlying spatial integration immediately poses two questions: (1) Where does this feedback originate from? and (2) Where is this feedback heading? We used fMRI recordings to answer these two questions in turn.

To reveal the source of the feedback, we evaluated how representations across visual cortex differ between stimuli that can or cannot be integrated across space (Fig. 2D). We reasoned that regions capable of exerting integration-related feedback should show stronger representations of spatiotemporally consistent inputs that demand integration, compared to inconsistent inputs that do not. Scene-selective areas in visual cortex are a strong contender for the source of this feedback, as they have been previously linked to the spatial integration of congruent scene information ^18,19^.

To test this assertion, we decoded the video stimuli from multi-voxel patterns in a set of three early visual cortex regions, V1, V2, and V3, and three scene-selective regions, the occipital place area (OPA), the medial place area (MPA), and the parahippocampal place area (PPA).

In a first step we decoded between the single videos, and found information only when video stimuli were shown in the hemifield contralateral to the region investigated (all *t*(35) > 3.75, *p* < 0.001; Fig. 2E). This implies that any stronger decoding for consistent, compared to inconsistent, video stimuli can only be driven by the interaction of ipsilateral and contralateral inputs, rather than by the ipsilateral input alone. On this interpretative backdrop, we next decoded congruent and incongruent video stimuli. Both were decodable in each of the six regions (all *t*(35) > 4.43, *p* < 0.001; Fig. 2F). Critically, congruent video stimuli were only better decoded than incongruent stimuli in the MPA (*t*(35) = 3.61, *p* < 0.001; Fig. 2F) and PPA (*t*(35) = 3.32, *p* = 0.002; Fig. 2F). In MPA and PPA, congruent video stimuli were also better decoded than contralateral single video stimuli (both *t*(35) > 3.2, *p* < 0.004). These results show that scene-selective cortex aggregates spatiotemporally consistent information across hemifields, suggesting PPA and MPA as likely generators of feedback signals guiding visual integration.

Finally, we determined where the alpha-frequency feedback is headed. We were particularly interested in whether integration-related feedback traverses the visual hierarchy up to the earliest stages of visual processing ^20,21^. To investigate this, we performed an EEG/fMRI fusion analysis ^22,23^ that directly links spectral representations in the EEG with spatial representations in the fMRI. To link representations across modalities, we first computed representational similarities between all videos using pair-wise decoding analyses and then correlated the similarities obtained from EEG alpha responses and fMRI activations across the six visual regions (Fig. 3A). Here, we focused on the crucial comparison of regional representations (fMRI) and alpha-frequency representations (EEG) between the congruent and the incongruent conditions. As feedforward inputs from the contralateral visual field are identical across both conditions, any stronger correspondence between regional representations and alpha-frequency representations in the consistent condition can unequivocally be attributed to feedback from higher-order systems, which have access to both ipsi- and contralateral input. We found that representations in the alpha band were more strongly related to representations in the congruent condition than in the incongruent condition in V1 (*t*(35) = 3.37, *p* = 0.001; Fig. 3B). A similar trend emerged in V2 (*t*(35) = 2.32, *p*_uncorrected_ = 0.025; Fig.3B) and V3 (*t*(35) = 2.15, *p*_uncorrected_ = 0.036; Fig. 3B) but not in scene-selective cortex (all *t*(35) < 1.67, *p* > 0.103; interaction: *F*(5, 235) = 10.17, *p* < 0.001; Fig. 3B). By contrast, no such correspondences were found between beta/gamma EEG responses and regional fMRI activations (see Fig. S4). The results of the fusion analysis show that when inputs are spatiotemporally consistent and demand integration into a unified percept, alpha-frequency feedback reaches down all the way to the earliest stages of visual processing in cortex.

**Fig. 3.**
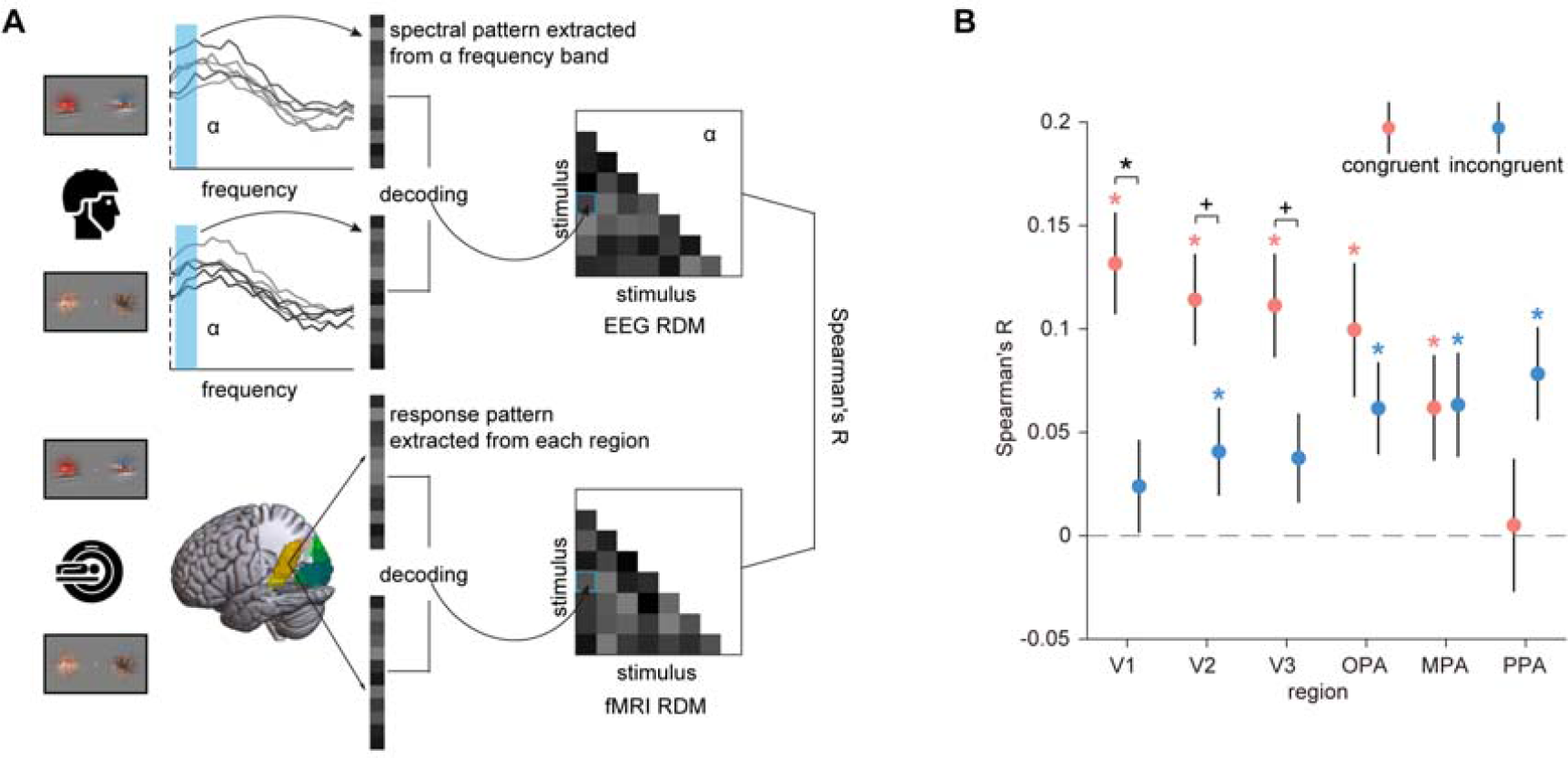
EEG-fMRI fusion analysis. **A)** For each condition, EEG representational dissimilarity matrices (RDMs) for each frequency band (α, β, γ) and fMRI RDMs for each region of interest (V1, V2, V3, OPA, MPA, PPA) were first obtained using pair-wise decoding analyses. To assess correspondences between spectral and regional representations, we calculated *Spearman*-correlations between the participant-specific EEG RDMs in each frequency band and the group-averaged fMRI RDMs in each region, separately for each condition. **B)** Results of EEG-fMRI fusion analysis in the alpha band. Representations in the alpha band corresponded more strongly with representations in V1 (with a similar trend in V2 and V3) when the videos were presented congruently, rather than incongruently. No correspondences were found between the beta and gamma bands and regional activity (see Fig. S4). Error bars represent standard errors. *: *p* < 0.05 (FDR-corrected), ^+^: *p* < 0.05 (uncorrected).

## Discussion

Our findings demonstrate that the spatial integration of natural inputs integral to mediating coherent perception is achieved by cortical feedback: Only when spatiotemporally coherent inputs demanded integration, stimulus-specific information was coded in feedback-related alpha activity. We further show that scene-selective areas in visual cortex interactively process information across visual space, highlighting them as likely sources of integration-related feedback. Finally, we reveal that integration-related alpha dynamics are linked to representations in early visual cortex, indicating that integration is accompanied by rhythmic feedback that traverses the whole cortical visual hierarchy from top to bottom. Together, our results promote an active conceptualization of the visual system, where concurrent feedforward and feedback information flows are critical for establishing coherent natural vision.

Our finding that feedback reaches all the way to initial stages of visual processing supports the emerging notion that early visual cortex receives various types of stimulus-specific feedback, such as during mental imagery ^24,25^, in cross-modal perception ^26,27^, and during the interpolation of missing contextual information ^28,29^. Further supporting the interpretation of such signals as long-range feedback, recent animal studies have found that contextual signals in V1 are substantially delayed in time, compared to feedforward processing ^30–32^. Such feedback processes may utilize the spatial resolution of V1 as a flexible sketchpad mechanism ^33,34^ for recreating detailed feature mappings that are inferred from global context.

Our fMRI data show that scene-selective areas in the anterior ventral temporal cortex (the MPA and the PPA) are the probable source of the feedback to early visual cortex. These regions exhibited stronger representations for spatiotemporally coherent stimuli placed in the two hemifields, despite a lack of sensitivity to the ipsilateral input alone. Scene-selective cortex is sensitive to the typical spatial configuration of scene stimuli ^18,19,35,36^, allowing it to create feedback signals that carry information about whether and how stimuli need to be integrated at lower levels of the visual hierarchy. To further support this notion, future studies should explicitly map cortico-cortical connectivity during natural perception.

Our results inform theories about the functional role of alpha rhythms in cortex. Alpha is often considered an idling rhythm ^37,38^, a neural correlate of active suppression ^39–41^, or a correlate of working memory maintenance ^42,43^. More recently, alpha rhythms were associated with an active role in cortical feedback processing ^12,14,44,45^. Our results highlight that alpha dynamics not only modulate feedforward processing, but encode stimulus-specific information. Our findings thus invite a new conceptualization of alpha dynamics, where alpha rhythms are recruited for the active transport of feedback information across the visual cortical hierarchy ^15,16,46,47^.

Our findings can be linked to theories of predictive processing that view neural information processing as a dynamic exchange of sensory feedforward signals and predictions stemming from higher-order areas of cortex ^5,6,46^. On this view, feedback signals arising during stimulus integration are conceptualized as predictions about sensory input derived from spatially and temporally coherent contralateral input. In our paradigm, feedback signals can be conceptualized as predictions for the contralateral input generated from the spatiotemporally consistent ipsilateral input. A challenge for predictive coding theories is that it requires a strict separation of the feedforward sensory input and the predictive feedback. Our results indicate a compelling solution through multiplexing of feedforward and feedback information in dedicated frequency-specific channels ^11,13,14,44^ in human cortex. It will be interesting to see whether similar frequency-specific correlates of predictive processing are unveiled in other brain systems in the future.

More generally, our results have general implications for understanding and modelling feedforward and feedback information flows in neural systems. Processes like stimulus integration that are classically conceptualized as solvable in pure feedforward cascades may be more dynamic than previously thought. Discoveries like ours arbitrate between competing theories that either stress the power of feedforward hierarchies ^3,4^ or emphasize the critical role of feedback processes ^5,6^. They further motivate novel approaches to computational modelling that capture the visual system’s abundant feedback connectivity ^48–50^.

Taken together, our results reveal feedback in the alpha frequency range across the visual hierarchy as a key mechanism for integrating spatiotemporally consistent information in natural vision. This strongly supports an active conceptualization of the visual system, where top-down projections are critical for the construction of coherent and unified visual percepts from fragmented sensory information.

## Methods

### Participants

Forty-eight healthy adults (gender: 12 M/36 F, age: 21.1 ± 3.8 years) participated in the EEG experiment, and another thirty-six (gender: 17 M/19 F, age: 27.3 ± 2.5 years) participated in the fMRI experiment. Sample size resulted from convenience sampling, with the goal of exceeding n = 34 in both experiments (i.e., exceeding 80% power for detecting a medium effect size of d = 0.5 in a two-sided *t*-test). All participants had normal or corrected-to-normal vision. They all signed informed consent before the experiment, and they were compensated for their time with partial course credit or cash. The EEG protocol was approved by the ethical committee of the Department of Psychology at the University of York, and the fMRI protocol was approved by the ethical committee of the Department of Psychology at Freie Universität Berlin. All experimental protocols were in accordance with the Declaration of Helsinki.

### Stimuli and design

The stimuli and design were identical for the EEG and fMRI experiments unless stated otherwise.

Eight short video clips (3 s each; Fig. 1A) depicting everyday situations (e.g., a train driving past the observer; a view of red mountains; a view of waves crashing on the coast; first-person perspective of walking along a forest path; an aerial view of a motorway toll station; a group of zebras grazing on a prairie; first-person perspective of skiing in the forest; first-person perspective of walking along a street) were used in the experiments. During the experiment, these original videos were presented on the screen through circular apertures right and left of central fixation. We manipulated four experimental conditions (right-only, left-only, congruent, and incongruent) by presenting the original videos in different ways (Fig. 1B). In the right- or left-only condition, we presented the videos through right or left aperture only. We also showed two matching segments from the same video in the congruent condition, while showed segments from two different videos in the incongruent condition, through both apertures. In the incongruent condition, the eight original videos were yoked into eight fixed pairs (see Fig. S1), and each video was always only shown with its paired video. Thus, there were a total of 32 unique video stimuli (8 for each condition). The diameter of each aperture was 6° visual angle, and the shortest distance between the stimulation and central fixation was 2.64° visual angle. The borders of the apertures were slightly smoothed. The central fixation dot subtended 0.44° visual angle.

The experiments were controlled through MATLAB and the Psychophysics Toolbox ^51,52^. In each trial, a fixation dot was first shown for 500 ms, after which a unique video stimulus was displayed for 3,000 ms. During the video stimulus playback, the color of the fixation changed periodically (every 200 ms) and turned either green or yellow at a single random point in the sequence (but never the first or last point). After every trial, a response screen prompted participants to report whether a green or yellow fixation dot was included in the sequence. Participants were instructed to keep central fixation during the sequence so they would be able to solve this task accurately. In both experiments, participants performed the color discrimination with high accuracy (EEG: 93.28 ± 1.65% correct; fMRI: 91.44 ± 1.37% correct), indicating that they indeed focused their attention on the central task. In the EEG experiment, the next trial started once the participant’s response was received. In the fMRI experiment, the response screen stayed on the screen for 1,500 ms, irrespective of participants’ response time. An example trial is shown in Fig. 1C.

In the EEG experiment, each of the 32 unique stimuli was presented 20 times, resulting in a total of 640 trials, which were presented in random order. In the fMRI experiment, participants performed 10 identical runs. In each run, each unique stimulus was presented twice, in random order. Across the 10 runs, this also resulted in a total of 640 trials.

### EEG recording and preprocessing

EEG signals were recorded using an ANT Waveguard 64-channel system and a TSMi REFA amplifier, with a sample rate of 1000 Hz. The electrodes were arranged according to the standard 10-10 system. EEG data preprocessing was performed using FieldTrip ^53^. The data were first band-stop filtered to remove 50Hz line noise and then band-pass filtered between 1 Hz and 100 Hz. The filtered data were epoched from -500 ms to 4,000 ms relative to the onset of the stimulus, re-referenced to the average over the entire head, downsampled to 250 Hz, and baseline corrected by subtracting the mean pre-stimulus signal for each trial. After that, noisy channels and trials were removed by visual inspection, and the removed channels were interpolated by the mean signals of their neighboring channels. Blinks and eye movement artifacts were removed using independent component analysis (ICA) and visual inspection of the resulting components.

### EEG power spectrum analysis

Spectral analysis was performed using Fieldtrip. Powerspectra were estimated between 8 and 70 Hz (from alpha to gamma range), from 0 to 3,000 ms (i.e., the period of stimulus presentation) on the preprocessed EEG data, separately for each trial and each channel. A single taper with a Hanning window was used for the alpha band (8–12 Hz, in steps of 1 Hz) and the beta band (13–30 Hz, in steps of 2 Hz), and the discrete prolate spheroidal sequences (DPSS) multitaper method with ±8 Hz smoothing was used for the gamma band (31–70 Hz, in steps of 2 Hz).

### EEG decoding analysis

To investigate whether the dynamic integration of information across the visual field is mediated by oscillatory activity, we performed multivariate decoding analysis using CoSMoMVPA ^54^ and LIBSVM ^55^. In this analysis, we decoded between the eight video stimuli using patterns of spectral power across channels, separately for each frequency band (alpha, beta, and gamma) and each condition. Specifically, for each frequency band, we extracted the power of the frequencies included in that band (e.g., 8–12 Hz for the alpha band) across all channels from the power spectra, and then used the resulting patterns across channels and frequencies to classify the eight video stimuli in each condition. For all classifications, we used linear support vector machine (SVM) classifiers to discriminate the eight stimuli in a ten-fold cross-validation scheme. For each classification, the data were allocated to 10 folds randomly, and then a SVM classifier was trained on data from 9 folds and tested on data from the left-out fold. The classification was done repeatedly until every fold was left out once, and accuracies were averaged across these repetitions. The amount of data in the training set was always balanced across stimuli. For each classification, a maximum of 144 trials (some trials were removed during preprocessing) were included in the training set (18 trials for each stimulus) and 16 trials were used for testing (2 trials for each stimulus). Before classification, principal component analysis (PCA) was applied to reduce the dimensionality of the data ^56^. Specifically, for each classification, PCA was performed on the training data, and the PCA solution was projected onto the testing data. For each PCA, we selected the set of components that explained 99% of the variance of the training data. As a result, we obtained decoding accuracies for each frequency band and each condition, which indicated how well the video stimuli were represented in frequency-specific neural activity. We first used a one-sample *t*-test to investigate whether the video stimuli could be decoded in each condition and each frequency band. We also performed a 2 condition (congruent, incongruent) × 3 frequency (alpha, beta, gamma) two-way ANOVA, and post-hoc paired *t*-tests (FDR-corrected across frequencies; *p*_corrected_ < 0.05) to compare the decoding differences between congruent and incongruent conditions separately for each frequency band. The comparisons of right- and left-only conditions were conducted using the same approaches. To track where the effects appeared across a continuous frequency space, we also decoded between the eight stimuli at each frequency from 8 to 70 Hz using a sliding window approach with a 5-frequency resolution (see Fig. S2).

### fMRI recording and processing

MRI data were acquired using a 3T Siemens Prisma scanner (Siemens, Erlangen, Germany) equipped with a 64-channel head coil. T2*-weighted BOLD images were obtained using a multiband gradient-echo echo-planar imaging (EPI) sequence with the following parameters: multiband factor = 3, TR = 1,500 ms, TE = 33 ms, FOV = 204 × 204 mm^2^, voxel size = 2.5 × 2.5 × 2.5 mm^3^, 70° flip angle, 57 slices, 10% interslice gap. Field maps were also obtained with a double-echo gradient echo field map sequence (TR = 545 ms, TE1/TE2 = 4.92 ms/7.38 ms) to correct for distortion in EPI. In addition, a high-resolution 3D T1-weighted image was collected for each participant (MPRAGE, TR = 1,900 ms, TE = 2.52 ms, TI = 900 ms, 256 × 256 matrix, 1 × 1 × 1 mm^3^ voxel, 176 slices).

MRI data were preprocessed using MATLAB and SPM12 (https://www.fil.ion.ucl.ac.uk/spm/). Functional data were first corrected for geometric distortion with the SPM FieldMap toolbox ^57^ and realigned for motion correction. In addition, individual participants’ structural images were coregistered to the mean realigned functional image, and transformation parameters to MNI standard space (as well as inverse transformation parameters) were estimated.

The GLMsingle Toolbox ^58^ was used to estimate the fMRI responses to the stimulus in each trial based on realigned fMRI data. To improve the accuracy of trial-wise beta estimations, a three-stage procedure was used, including identifying an optimal hemodynamic response function (HRF) for each voxel from a library of 20 HRFs, denoising data-driven nuisance components identified by cross-validated PCA, and applying fractional ridge regression to regularize the beta estimation on a single voxel basis. The resulting single-trial betas were used for further decoding analyses.

### fMRI regions of interest definition

fMRI analyses were focused on six regions of interest (ROIs). We defined three scene-selective areas – occipital place area (OPA; also termed transverse occipital sulcus ^59,60^), medial place area (MPA; also termed retrosplenial cortex ^61,62^), and parahippocampal place area (PPA ^63^) – from a group functional atlas ^64^; and three early visual areas – V1, V2, and V3 – from a probabilistic functional atlas ^65^. All ROIs were defined in MNI space and separately for each hemisphere, and then transformed into individual-participant space using the inverse normalization parameters estimated during preprocessing.

### fMRI ROI decoding analysis

To investigate how the video stimuli were processed in different visual regions, we performed multivariate decoding analysis using CoSMoMVPA and LIBSVM. For each ROI, we used the beta values across all voxels included in the region to decode between the eight video stimuli, separately for each condition. Leave-one-run-out cross-validation and PCA were used to conduct SVM classifications. For each classification, there were 144 trials (18 for each stimulus) in the training set and 16 trials (2 for each stimulus) in the testing set. For each participant, we obtained a 4 condition × 12 ROI (6 ROIs by 2 hemispheres) decoding matrix. Results were averaged across hemispheres, as we consistently found no significant inter-hemispheric differences (condition × hemisphere and condition × region × hemisphere interaction effects) in a 2 condition (congruent, incongruent) × 6 region (V1, V2, V3, PPA, OPA, MPA) × 2 hemisphere (left, right) three-way ANOVA test. We first tested whether the video stimuli were decodable in each condition and each region using one-sample *t*-tests (FDR-corrected across regions; *p*_corrected_ < 0.05). To further investigate the integration effect, we used paired *t*-tests to compare the decoding difference between congruent and incongruent conditions in different regions. The same approaches were also used to compare the two single video conditions.

### EEG-fMRI fusion with representational similarity analysis

To investigate the relationship between the frequency-specific effects obtained in the EEG and the spatial mapping obtained in the fMRI, we performed EEG-fMRI fusion analysis ^22,23^. This analysis could be used to compare neural representations of stimuli as characterized by EEG and fMRI data to reveal how the representations correspond across space and time.

Specifically, we first calculated representational dissimilarity matrices (RDMs) using pair-wise decoding analysis for EEG and fMRI data, respectively. For the EEG power spectra, in each frequency band, we decoded between each pair of eight video stimuli using the oscillatory power of the frequencies included in the frequency band, separately for each condition; for the fMRI data, in each ROI, we classified each pair of eight stimuli using the response patterns of the region, separately for each condition. Decoding parameters were otherwise identical to the 8-way decoding analyses (see above). In each condition, we obtained a participant-specific EEG RDM (8 stimuli × 8 stimuli) in each frequency band and a participant-specific fMRI RDM (8 stimuli × 8 stimuli) in each ROI. Next, we calculated the similarity between EEG and fMRI RDMs for each condition; this was done by correlating all lower off-diagonal entries between the EEG and fMRI RDMs (the diagonal was always left out). To increase the signal-to-noise ratio, we first averaged fMRI RDMs across participants and then calculated the *Spearman*-correlation between the averaged fMRI RDM for each ROI with the participant-specific EEG RDM for each frequency. As a result, we obtained a 4 condition × 3 frequency × 12 ROI fusion matrix for each EEG participant. The results were averaged across regions, as no condition × hemisphere, no condition × region × hemisphere, and no condition × frequency × hemisphere interaction effects were found. We first used one-sample *t*-tests to test the fusion effect in each condition (FDR-corrected across regions; *p*_corrected_ < 0.05) and each frequency-region combination, and then used a 2 condition × 3 frequency × 6 region three-way ANOVA to compare the frequency-region correspondence between congruent and incongruent conditions. As we found a significant condition × frequency × region interaction effect, we further performed a 2 condition × 6 region ANOVA and paired *t*-tests (FDR-corrected across regions; *p*_corrected_ < 0.05) to compare frequency-region correspondence between congruent and incongruent conditions separately for each frequency. The comparisons of right- and left-only conditions were performed using the same approach (see Fig. S4).

## Acknowledgements

L.C. is supported by a PhD stipend from the China Scholarship Council (CSC). R.M.C is supported by the Deutsche Forschungsgemeinschaft (DFG; CI241/1-1, CI241/3-1, CI241/7-1) and by a European Research Council (ERC) starting grant (ERC-2018-STG 803370). D.K. is supported by the Deutsche Forschungsgemeinschaft (DFG; SFB/TRR135 – INST162/567-1), a European Research Council (ERC) starting grant (ERC-2022-STG 101076057), and “The Adaptive Mind”, funded by the Excellence Program of the Hessian Ministry of Higher Education, Science, Research and Art. The authors thank Daniela Marinova and Alex Carter for help in EEG data acquisition. The authors would also like to thank the HPC Service of ZEDAT, Freie Universität Berlin^66^, for computing time.

## Author contributions

Conceptualization, L.C. and D.K.; Methodology, L.C. and D.K.; Software, L.C. and D.K.; Formal analysis, L.C.; Investigation, L.C. and D.K.; Resources, L.C. and D.K.; Data curation, L.C.; Writing **–** original draft, L.C. and D.K.; Writing **–** review & editing, L.C., R.M.C., and D.K.; Visualization, L.C.; Supervision, R.M.C. and D.K.; Project administration, R.M.C. and D.K.; Funding acquisition, R.M.C. and D.K.

## Declaration of interests

The authors declare no competing interests.

## Supplementary Information (SI)

**Fig. S1.**
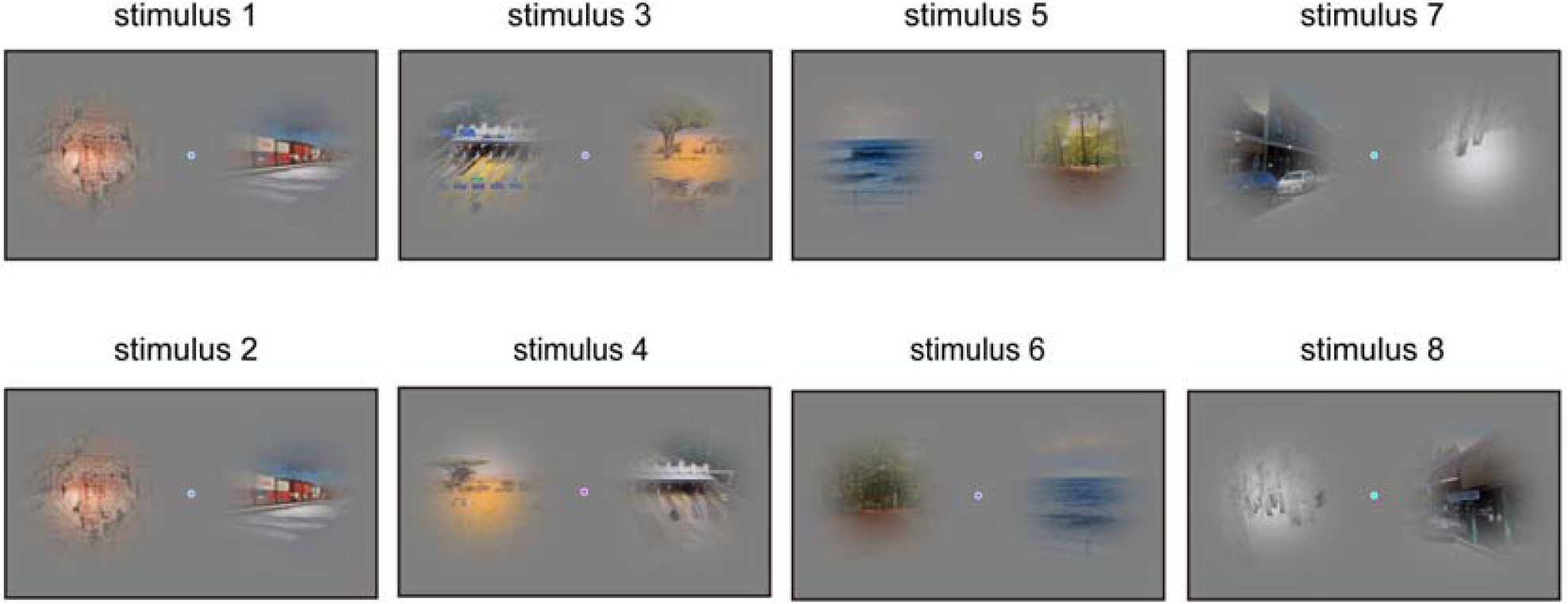
Incongruent stimuli.

**Fig. S2.**
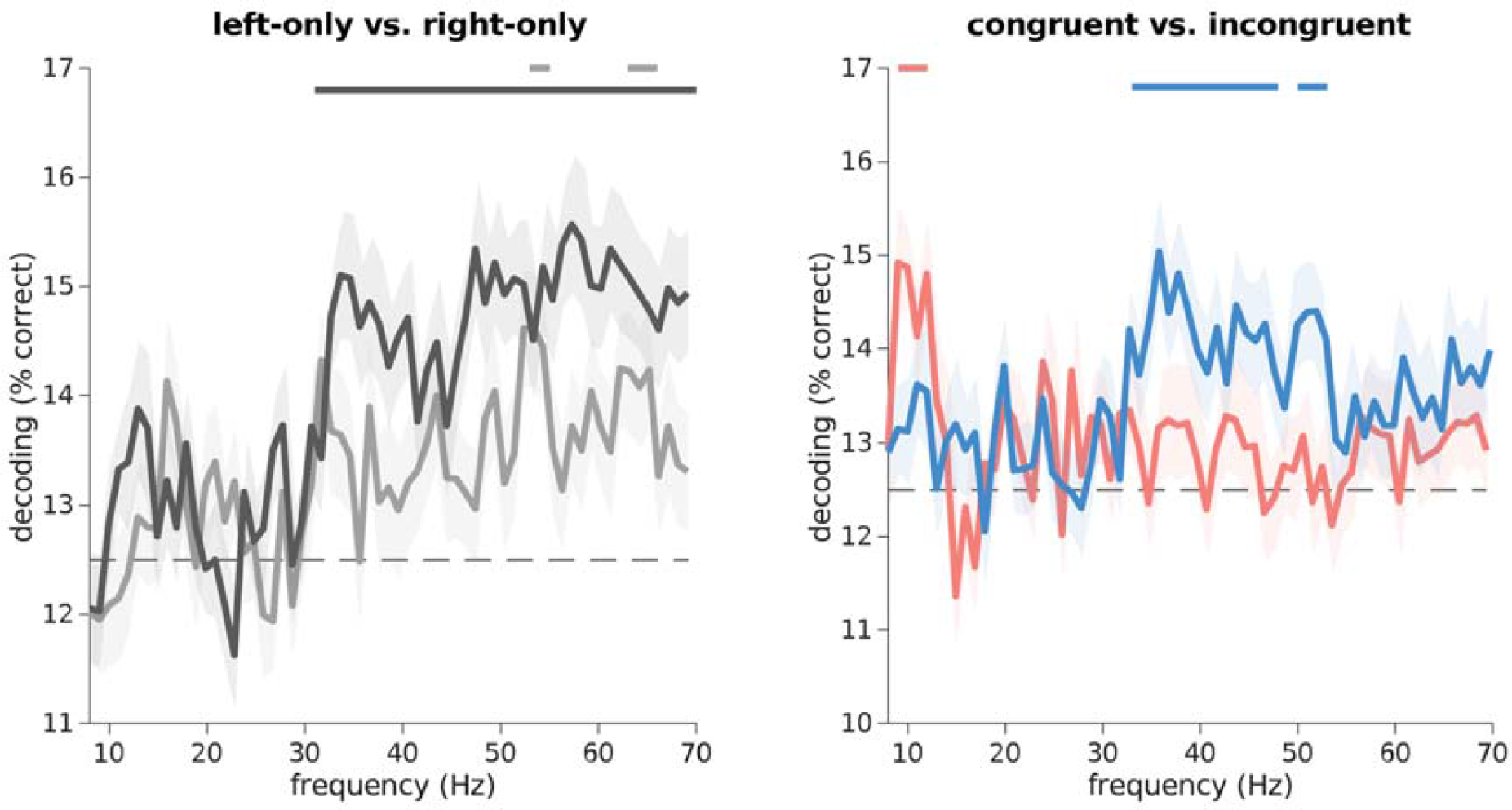
EEG frequency-resolved decoding analysis on oscillatory power patterns using sliding window (8–70 Hz). We decoded between the eight video stimuli at each frequency from 8 to 70 Hz using a sliding window approach with a 5-frequency resolution, separately for each condition. The incongruent and single video stimuli were decodable from the gamma frequency band, whereas congruent videos were decodable from the alpha frequency band. Line markers denote significant above-chance decoding (*p* < 0.05; FDR-corrected).

**Fig. S3.**
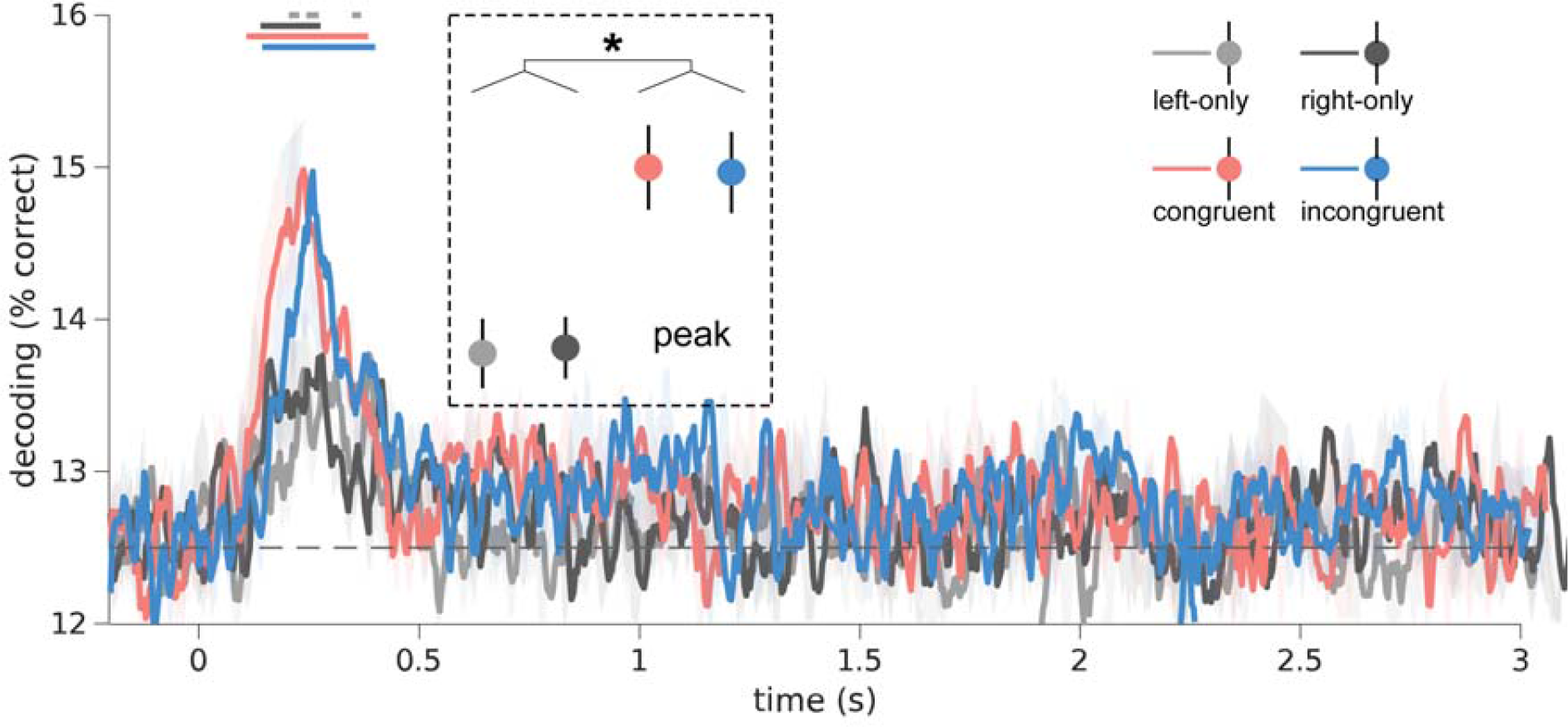
EEG time-resolved decoding on evoked response patterns (-0.2–3 s). We performed decoding analysis on time-resolved broadband responses across channels to discriminate the eight video stimuli at each time from -200 ms to 3,000 ms relative to the onset of the stimulus, separately for each condition. The obtained decoding timeseries for each condition were smoothed by the moving average algorithm (6 time points). We extracted the peak decoding accuracy for each condition and then compared the decoding difference between conditions using paired *t*-tests. The results revealed a sustained representation of the video stimuli across the first second of processing, with stronger peak responses to two video conditions (congruent/incongruent) than to single video conditions (right-/left-only), but no differences between the congruent and incongruent conditions. Line markers denote significant above-chance decoding. *: *p* < 0.05 (FDR-corrected).

**Fig. S4.**
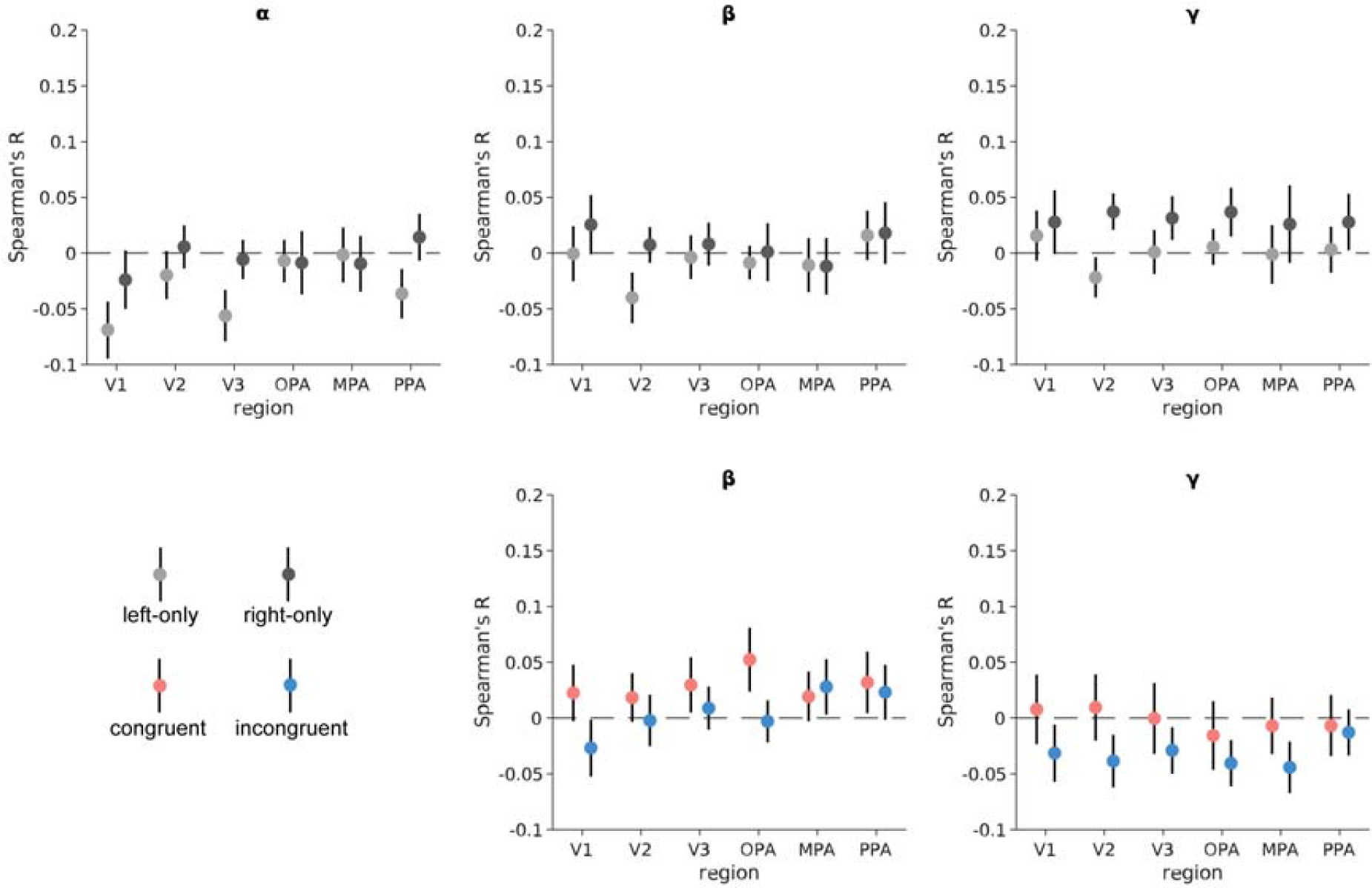
EEG-fMRI fusion analysis separately for each frequency band. For each condition, EEG representational dissimilarity matrices (RDMs) for each frequency band (α, β, γ) and fMRI RDMs for each region of interest (V1, V2, V3, OPA, MPA, PPA) were first obtained using pair-wise decoding analyses. To assess correspondences between spectral and regional representations, we calculated Spearman-correlations between the participant-specific EEG RDMs in each frequency band and the group-averaged fMRI RDMs in each region, separately for each condition. For the right- and left-only conditions, there was no significant correspondence between EEG responses in each frequency band (α, β, γ) and fMRI activations in each region (V1, V2, V3, OPA, MPA, PPA). For the congruent and incongruent conditions, there were no significant correspondences between beta/gamma responses and fMRI activations. Error bars represent standard errors.

## Notes

### Competing Interest Statement

The authors have declared no competing interest.

### Summary of Updates

update manuscript

